# Engineering Memory T Cells as a platform for Long-Term Enzyme Replacement Therapy in Lysosomal Storage Disorders

**DOI:** 10.1101/2024.04.23.590790

**Authors:** Kanut Laoharawee, Evan W Kleinboehl, Jacob D Jensen, Joseph J Peterson, Nicholas J Slipek, Bryce J Wick, Matthew J Johnson, Beau R Webber, Branden S Moriarity

## Abstract

Enzymopathy disorders are the result of missing or defective enzymes. Amongst these enzymopathies, mucopolysaccharidosis type I, is a rare genetic lysosomal storage disorder caused by mutations in the gene encoding alpha-L-iduronidase (IDUA), ultimately causes toxic build-up of glycosaminoglycans (GAGs). There is currently no cure and standard treatments provide insufficient relief to the skeletal structure and central nervous system (CNS). Human memory T cells (Tm) migrate throughout the body’s tissues and can persist for years, making them an attractive approach for cellular-based, systemic enzyme replacement therapy. Here, we tested genetically engineered, IDUA-expressing Tm as a cellular therapy in an immunodeficient mouse model of MPS I. Our results demonstrate that a single dose of engineered Tm leads to detectable IDUA enzyme levels in the blood for up to 22 weeks and reduced urinary GAG excretion. Furthermore, engineered Tm take up residence in nearly all tested tissues, producing IDUA and leading to metabolic correction of GAG levels in the heart, lung, liver, spleen, kidney, bone marrow, and the CNS. Our study indicates that genetically engineered Tm holds great promise as a platform for cellular-based enzyme replacement therapy for the treatment of mucopolysaccharidosis type I and potentially many other enzymopathies and protein deficiencies.

## Introduction

Enzymopathy disorders resulting from various genetic mutations such as Mucopolysaccharidosis type I (MPS I), also called Hurler-Scheie Syndrome, is a rare lysosomal storage disorder (LSD) characterized by insufficient alpha-L-iduronidase (IDUA) enzyme activity due to mutation of the IDUA gene.^1,2^ IDUA is essential for the multi-step degradation pathway of glycosaminoglycans (GAGs). IDUA deficiency leads to harmful accumulation of un-degraded GAGs within the lysosome, resulting in a progressive pathophysiology involving all major body systems.^2,3^ Symptoms include musculoskeletal dysfunction, skeletal deformation, heart valve disease, central nervous system (CNS) dysfunction, neurocognitive dysfunction, corneal clouding, and a significantly decreased lifespan.^2,4^ To date, there is no cure for MPS I, and current therapies are limited to either enzyme replacement therapy (ERT), in which patients are periodically given infusions of recombinant IDUA, and/or an allogeneic haematopoietic stem cell transplantation (HSCT).^1,5^ Efficacy of these treatments is dependent on the process of systemic cross-correction, in which the patient’s own cells are able to endocytose functional extracellular IDUA via mannose-6-phosphate (M-6-P) receptor and the best results occur when treatment is started at the earliest possible age.^1,5,6^

ERT involves repeated peripheral infusions of recombinant IDUA (Laronidase/Aldurazyme) enzyme.^5^ Unfortunately, due to a short half-life of 1.5-3.6 hours, the enzyme concentration is not sustained long enough for a lasting therapy.^5^ In addition, ERT does not improve neurocognitive deficits in the affected individuals due to the limited ability of the enzyme to cross the blood-brain barrier (BBB).^5,7^ ERT is also very costly (> $250,000 per year) and requires frequent, time-consuming visits to the clinic, which have a significant negative impact on the patient’s quality of life and healthcare resources. ^6^ Allogeneic HSCT is the current standard of care for severe cases of MPS I because it offers a one-time procedure that can provide lifetime supplementation of IDUA.^6,8^ HSCT also allows IDUA to be systemically delivered, although this can take up to a year post-transplant before effects are observed.^6,9–11^ However, allogeneic HSCT comes with several critical drawbacks, including scarce HLA-matched donors, risk of complications and death associated with graft-vs-host disease (GVHD), low engraftment rates, and typically results in insufficient enzyme levels, especially within the CNS.^3,6,9,12,13^

T cells are adaptive immune cells derived from lymphoid-lineage progenitors of hematopoietic stem cells in the bone marrow. They are one of the most abundant leukocytes in the blood and can be easily isolated from peripheral blood without need for bone marrow mobilization. In addition, T cells have been shown to cross the BBB. ^14,15^ Two main subsets of T cells exist, CD4+ helper T cells, which have numerous roles in regulating immune responses, and CD8+ cytotoxic T cells, which have the ability to directly kill other cells. Upon activation via T cell receptor (TCR) signaling, T cells can differentiate further into a variety of subtypes, including CD45RO+ memory T cells (Tm), which are capable of maintaining a population within the body for decades.^16–18^ In this study, we engineer CD4+ CD45RO+ memory helper T cells as IDUA enzyme producing factories in order to take advantage of the mobility and persistence of these cells. This approach using memory CD4+ T cells minimizes graft-versus-host disease (GVHD) in our animal model rather than using total CD3+ cells.^19^ The accessibility, mobility, and distribution of Tm cells throughout the body combined with the potential to generate cells that persist for long periods of time make these cells an attractive cellular chassis for use in cellular gene therapies. Future therapies using autologous genetically modified T cells to secrete an enzyme of interest offers a flexible avenue for therapy while eliminating the issues associated with donor matching and rejection in allogeneic HSCT.^16,17,20,21^

Driven by the success of FDA approved CAR-T therapies, the engineered T cell field has matured significantly, leading to mass optimization of manufacturing techniques and development of novel tools for T cell-based therapeutics. Precision gene modification of primary human T cells has thus far focused largely on expressing chimeric antigen receptors (CAR-T) as anti-cancer therapeutics, yet this technology has applications outside the field of cancer immunotherapy, and the tools that have been developed by that field can easily be leveraged for novel T cell-based protein delivery systems. ^21–27^

The CRISPR-Cas9 system mediates precise insertion of transgenes by the induction of a targeted double-stranded DNA break (DSB) along with a DNA template for homology directed repair (HDR).^21,28,29^ Some recent studies have reported results with T cells engineered to produce and secrete various proteins via transposon or lentiviral methods.^27,30^ However, to date, no therapies have explored T cells precision-engineered at a safe harbor locus as an enzyme delivery platform for enzymopathies.

Here, we describe a novel process for CRISPR-Cas9 mediated, site-specific, insertion of a therapeutic human *IDUA* gene cassette at the *AAVS1* locus of human Tm cells. We demonstrate that CRISPR-Cas9 engineered Tm cells can express large quantities of enzymatically active IDUA. Furthermore, we demonstrate that a single dose of engineered Tm cells can persist in an immunodeficient mouse model of MPS I (NSG-MPS I)^31^ and secrete enzymatically active IDUA for up to 22 weeks. Persistence of the enzyme within the model resulted in systemic cross correction in various vital organs, reduced tissue pathology, and partially normalized urine and tissue GAG contents. Finally, we demonstrate that persistence of IDUA in the system slows the manifestation of neurocognitive deficiency of NSG-MPS I mice.

## Results

### Human Tm cells can be engineered to express high levels of IDUA

To test whether Tm cells can be engineered to express and excrete therapeutic IDUA protein, we utilized CRISPR-Cas9 combined with rAAV6 to deliver a DNA donor template for site-specific integration of an MND-IDUA-RQR8 **(Figure 1A)** construct at the *AAVS1* locus of CD4+ Tm cells (**Supplementary** Figure 1). RQR8 contains epitopes from CD34 and CD20 that can be used as a cell surface reporter and a target for GMP immunomagnetic enrichment, or as a safety mechanism allowing targeted depletion of the engineered cells using Rituximab (anti-CD20 monoclonal antibody).^33,34^ To this end, Tm cells were electroporated with Cas9 mRNA and sgRNA targeting the *AAVS1* locus and transduced with rAAV6 carrying the DNA donor molecule for HDR at the *AAVS1* locus. We observed up to ∼50% RQR8+ Tm cells at day 8 post engineering, while no RQR8+ cells were detected in transduction-only control cells **(Figure 1B)**. We also observed high levels of IDUA enzymatic activity in the medium containing engineered Tm cells, and negligible IDUA activity in the medium containing non-engineered control Tm cells **(Figure 1C)**. These data demonstrate that CRISPR-Cas9 gene editing along with rAAV6 as a DNA template donor can mediate efficient transgene integration at the *AAVS1* locus in primary human Tm cells, leading to robust production and excretion of functional IDUA enzyme *in vitro*.

**Figure 1.**
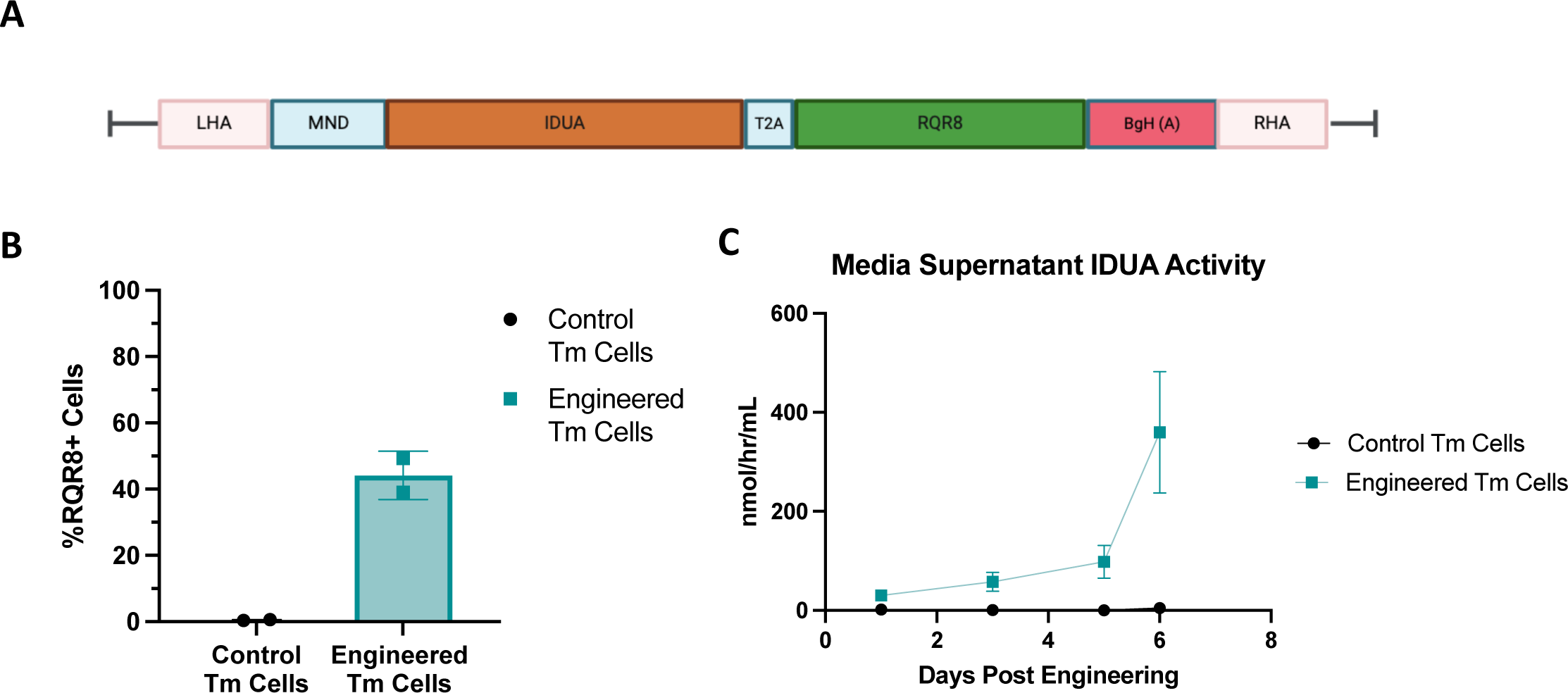
Efficient transgene insertion at the AAVS1 locus of human Tm cells using CRISPR-Cas9 and rAAV. (A) Schematic of rAAV IDUA expression cassette encoding homology arms targeting the *AAVS1* locus andMND promoter driving expression of IDUA-T2A-RQR8. (B) Quantification of RQR8 expression via flow cytometry in engineered Tm and non-engineered Tm cells. (C) Line graph depicting IDUA activity over time in the supernatant of engineered and control Tm cells (N=2 independent donors).

### Engraftment of engineered IDUA-expressing Tm cells demonstrate systemic correction of IDUA levels and GAG contents in a 10-week short-term study

We initiated a 10-week *in vivo* short-term study using an NSG model of Hurler syndrome (NSG-MPS I) to assess the efficacy of short-term treatment involving engineered IDUA-expressing Tm cells. This study consisted of three groups: IDUA+/- NSG mice as positive controls, untreated IDUA-/- NSG mice as MPS I controls, and IDUA-/- NSG mice treated with with IDUA engineered Tm cells;group). Both control groups were IP injected with vehicle (PBS). The Tm-treated group received 10 million bulk engineered Tm cells per animal. Human leukocyte marker, hCD45RO was used to evaluate circulating Tm cells in the periphery over time. The percent of hCD45RO+ leukocytes in peripheral blood increased throughout the first six weeks of the study and then diminished until week ten **(Figure 2A)**. Similarly, plasma IDUA activity in the Tm-treated NSG-MPS I mice increased two weeks post injection and continued to increase until week six, before declining to a level between untreated mutant and heterozygous controls by week 10 **(Figure 2B)**. The presence of hCD45RO+ cells and significant IDUA plasma activity indicated successful *in vivo* secretion of IDUA by engineered Tm cells.

**Figure 2.**
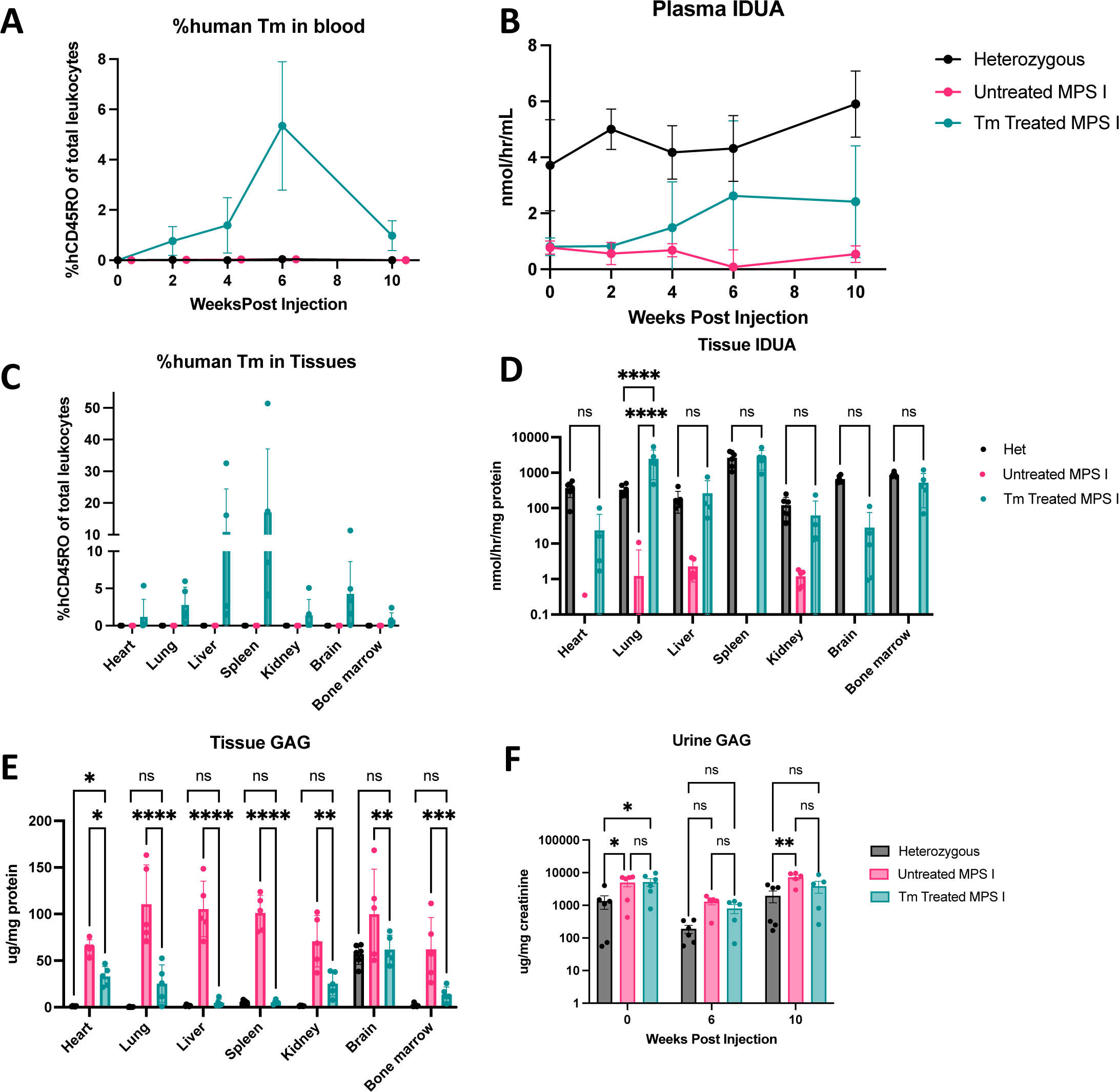
Short term efficacy of engineered Tm cells to secrete IDUA and ameliorate waste substrates *in vivo*. (A) Flow cytometry results of human CD45RO+ cells in total peripheral blood leukocytes in mice over time. (B) Fluorometric assay of murine plasma IDUA activity over a 10 week period. (C) Flow cytometry results of human CD45RO+ cells within organ tissues. (D) Fluorometric assay of tissue lysate IDUA activity levels at week 10 in multiple organs. (E) Tissue GAG content of tissue lysates at week 10 post treatment. (F) Urine GAG content at week 0, 6, and 10 post Tm cell treatment. All figures: Heterozygous (black), Untreated MPS I (red), and Tm-treated MPS I (blue) NSG mice. N=5 mice each cohort. Statistics analysis: Two-way ANOVA *p<0.05, **p<0.01, ***p<0.001, ****p,0.0001. (F) Error bars represent mean with SD.

At week 10, mice were sacrificed and perfused to clear the organs and tissues of blood. The heart, lung, liver, spleen, kidney, spinal cord, brain, and bone marrow tissues were collected and analyzed for the presence of engineered CD45RO+ Tm cells, enzymatically active IDUA, and GAG content. In the Tm-treated MPS I group, we observed a range of memory T cells within total leukocytes from the organs (1-60%) **(Figure 2C)**. As expected, we did not detect human Tm cells in the blood of control heterozygous or untreated NSG-MPS I mice. Within treated mice, the spleen contained the highest percentage of Tm cells compared to other organs. Excitingly, Tm cells were found in the brain and bone marrow, while minimal Tm cells resided in the heart and kidneys **(Figure 2C)**.

Tissue lysates were used to measure IDUA activity in each organ. As anticipated, we detected a significant increase in IDUA enzymatic activity in all tested organs of the Tm-treated mice **(Figure 2D)**. Aside from the heart and brain, all other organs showed comparable IDUA activity levels to the heterozygous control group. Interestingly, IDUA activity in the lungs of the Tm-treated mice surpassed that of the heterozygote controls **(Figure 2D)**. Although no Tm cells were observed in the heart or kidney, their IDUA levels were significantly higher than in untreated animals, suggesting potential absorption of IDUA enzyme from circulation. This data indicates that engineered Tm cells may supply IDUA both locally and systemically.

GAG content was assessed in the same tissue lysates. Compared to heterozygous mice, untreated MPS I animals showed significantly elevated GAG contents in all tested organs except spinal cord tissues **(Figure 2E)**. However, Tm-treated MPS I animals demonstrated normalized GAG content in the lung, liver, spleen, brain, and bone marrow compared to untreated MPS I animals **(Figure 2E)**. This data suggests that IDUA enzyme secreted by engineered Tm cells leads to substantial reductions in tissue GAG contents in these organs.

Previous studies have reported elevated GAGs in the urine of NSG-MPS I mice as a hallmark of MPS I.^31^ Studies also report that sustained enzyme levels in the system lead to reduced urine GAGs over time.^35,36^ Indeed, we observed elevated urine GAGs in the NSG-MPS I mice as early as three weeks of age prior to treatment **(Figure 2F, week 0-time point)** and untreated NSG-MPS I mice showed continued elevation in urine GAG content over time, whereas a slight reduction in urine GAGs was observed in Tm-treated NSG-MPS I mice at week ten post-engraftment compared to untreated MPS I **(Figure 2F).**

### Engineered IDUA-expressing Tm cell treatment allows for long-term expression and secretion of IDUA

To evaluate the long-term efficacy of IDUA-expressing Tm cell treatment, we conducted a study comprising three experimental groups, each with 12 mice: heterozygous, NSG-MPS I, and Tm-treated NSG-MPS I mice. The treated group received an infusion of 10 million bulk engineered Tm cells per mouse. We observed human leukocytes in mice infused with the engineered Tm cells increase through week 4 and steadily decrease after week 10 in the peripheral blood **(Figure 3A)**. Additionally, although there was a reduction in Tm cells over time, they persisted for at least 22 weeks in the peripheral blood post-infusion **(Figure 3A)**. Furthermore, the presence of Tm cells coincided with an elevation of IDUA activity in the plasma of these mice as early as week four, surpassing heterozygous activity levels at week six. IDUA activity in the plasma began to decrease after week 6, though it remained above the IDUA level of untreated mice throughout the entire study **(Figure 3B)**. Flow cytometry analysis performed on peripheral blood indicated that the percent of RQR8 positive Tm cells was maintained throughout the study, suggesting that the engineering process did not adversely affect the long-term survival of the Tm cells **(Figure 3C)**. These data are in line with our short term 10 week study and demonstrate that engineered Tm cells are able to persist in the NSG mice and deliver IDUA systematically for up to 22 weeks.

**Figure 3.**
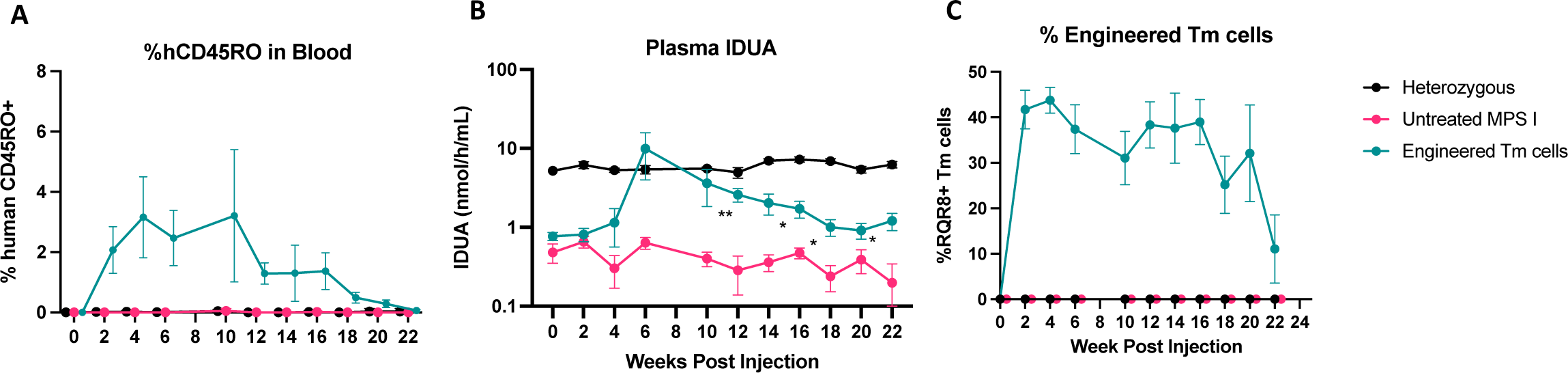
Engineered Tm cells persist and continually produce IDUA within NSG mice for up to 22 weeks. (A) Flow cytometry results of human CD45RO cells showing total leukocytes in the peripheral blood of Tm-treated NSG-MPS I mice. (B) Graph of fluorometric assay of murine plasma IDUA activity over 22 weeks from treated and control cohorts of mice. (C) Graph of RQR8 stained cells within the human CD45RO+ population of peripheral blood leukocytes as measured by flow cytometry. All figures: Heterozygous (black), and Tm-treated MPS I (blue) NSG mice. n=12 mice in each cohort. Statistical Analysis: Mixed Effects Analysis (A-C) *p<0.05, **p<0.01.

### The presence of IDUA expressing Tm cells in tissues correlates with a reduction of GAGs and lysosome accumulation in engineered Tm-treated NSG-MPS I mice

To determine the presence of the Tm cells residing in tissues at the end of the 22-week study, mice were sacrificed, perfused, and the heart, lung, liver, spleen, kidney, spinal cord, brain, and bone marrow tissues were collected to determine the presence of engineered Tm cells. Flow cytometry was used to identify human CD45RO+, CD3+ cells residing in the organs. Notably, we found fewer Tm cells in all tissues at 22 weeks (**Figure 4A**) compared to results in our short term 10 week study **(Figure 2C).** The bone marrow again had the largest proportion of Tm cells at an average of 1.1% of total cells, while the heart tissue exhibited the lowest concentration of identifiable Tm cells. Additionally, the CNS had detectable Tm cells at 22 weeks, indicating their potential to deliver IDUA to the CNS long term. As expected, we did not observe human cells in the untreated MPS I and the heterozygous mice.

**Figure 4.**
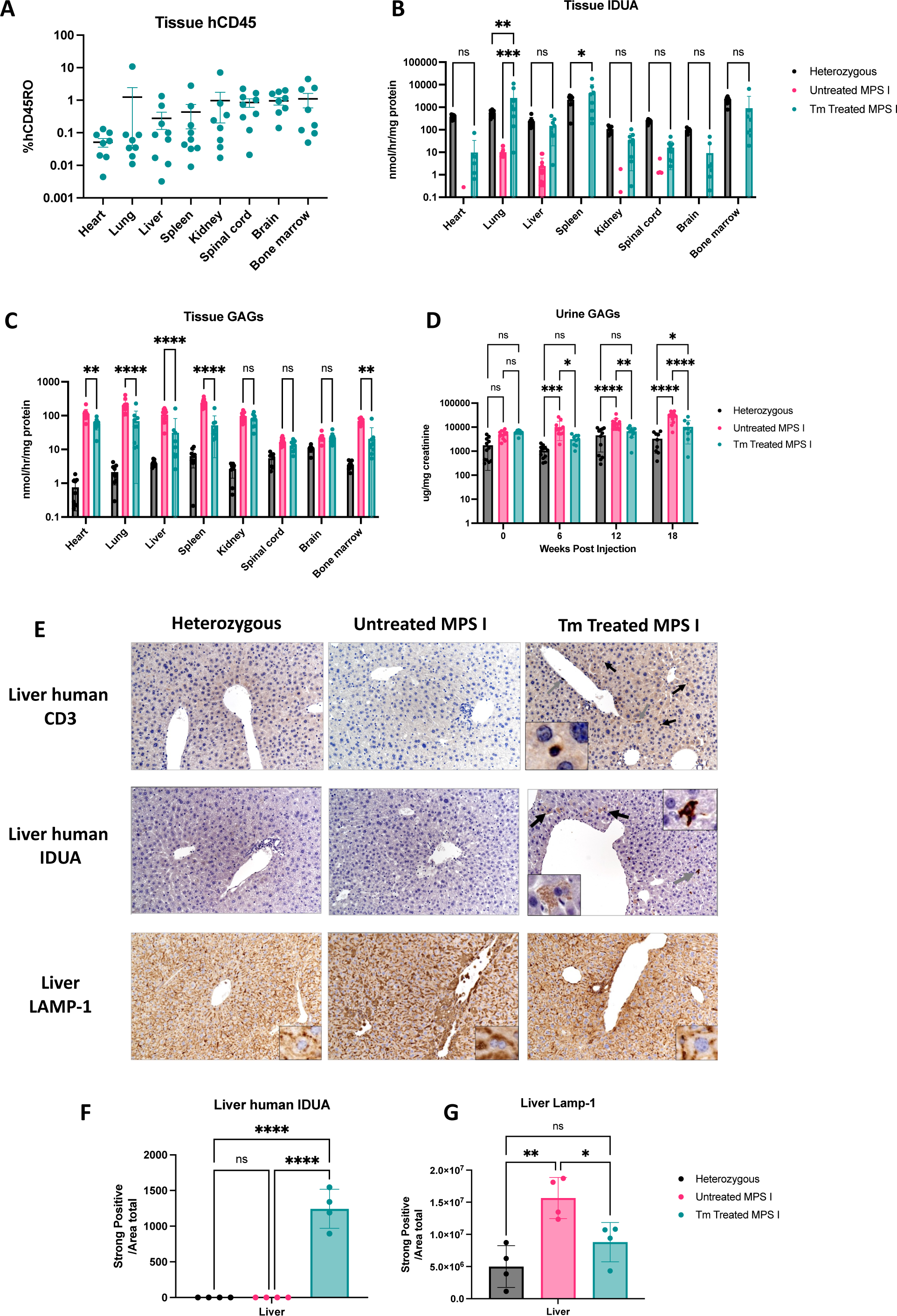
Engineered Tm cells provide IDUA systemically and reduce GAG accumulations across multiple tissues, ameliorating histopathological hallmarks of MPS I. (A) Flow cytometry results of human CD45RO within organs of Tm-treated NSG-MPS I mice. (B) Fluorometric assay of tissue lysate IDUA activity levels at week 22 post human Tm-treatment. (C) GAG content of tissue lysates at week 22 post human Tm cell treatment. (D) Urine GAG content at week 0, 6, 12, and 18 weeks post Tm cell treatment. (A-D) Heterozygous (black), NSG-MPS I (red), and Tm-treated NSG-MPS I (blue) NSG mice (n=12 mice each cohort). (E) Representative histological stained images of human CD3 (top), human IDUA (middle) and LAMP-1 (bottom) of mouse cohorts. (F) Quantification of human IDUA immunohistochemistry within liver samples (n=4 each cohort). (G) Quantification of LAMP-1 immunohistochemistry within liver samples (n=4 each cohort). Statistical Analysis: Two-way ANOVA (A-D), One-way ANOVA (F-G) *p<0.05, **p<0.01, ***p<0.001, ****p,0.0001.

Next, we evaluated the engineered cells’ efficacy in delivering IDUA locally and systemically to reduce MPS I hallmarks. Organ tissue lysates were used to measure local IDUA activity. Remarkably, there was no significant difference in tissue IDUA activity in the lung, liver, spleen, and bone marrow between engineered Tm-treated NSG-MPS I mice and heterozygous mice **(Figure 4B)**. All other organs from engineered Tm-treated NSG-MPS I mice had measurable IDUA activity levels present within the tissue lysate, albeit at levels lower than observed in heterozygous mice. These data indicate that the engineered Tm cells effectively deliver IDUA locally and systemically throughout treated mice for a minimum of 22 weeks. Next, the same tissue lysates from each organ were used for GAG assay to assess GAG contents. Notably, GAG contents of these lysates were significantly reduced in all tested organs, except the spinal cord **(Figure 4C)**. This indicates that IDUA secreted by engineered Tm cells can be taken up by host cells for cross correction and leads to restoration of the GAG degradation process.

We assessed urine GAG levels over the course of 18-week post infusion of engineered Tm cells. Despite the low level of IDUA in the plasma throughout the course of the study **(Figure 3B),** we observed normalization of urine GAGs in the treated mice at weeks 6 and 12, while urine GAGs of untreated MPS I animals continued to increase over time **(Figure 4D)**. Notably, the reduction in plasma IDUA activity levels of the Tm-treated mice after week 6 **(Figure 3B)**, coincided with an increase in urine GAGs over the same period **(Figure 4D)**.

At the 22-week endpoint, we evaluated a subset of liver samples from each group using immunohistochemical staining for human CD3, human IDUA, and LAMP-1 (lysosomal associated membrane protein 1). Increases in LAMP-1 are used as a diagnostic marker for lysosomal storage disorders, such as MPS I.^37,38^ Human CD3 staining within heterozygous and untreated mice showed no notable positive staining. The treated MPS I group had rare CD3 positive cells with lymphocyte morphology scattered throughout the sinusoids and portal triads (black arrows and insets) **(Figure 4E)**. Human IDUA staining within heterozygous and untreated NSG-MPS I mice showed no positive staining **(Figure 4E)**. Tm-treated NSG-MPS I mice showed rare, predominantly centrilobular regions with clusters of hepatocytes with cytoplasmic immunopositivity (black arrows). In other sections, there are scattered individual lymphocytes with strong cytoplasmic immunopositivity for human IDUA (black arrows) (**Figure 4E**). Quantification of staining in these liver samples from each group indicated no presence of human IDUA within heterozygous and NSG-MPS I mice while the Tm-treated NSG-MPS I mice had strong, significant immunopositivity **(Figure 4F)**. LAMP-1 stained samples from heterozygous mice contained fine granular, medium cytoplasmic immunopositivity at the edge and perinuclear zone of the hepatocytes. Untreated MPS I samples had strong cytoplasmic immunopositivity along the edges of the hepatocytes. Distribution of immunopositivity in Tm-treated NSG-MPS I mice was more similar to heterozygous mice; however, there was a slightly coarse granular and stronger cytoplasmic immunopositivity **(Figure 4E)**. Quantification of LAMP-1 staining within these samples indicated a significant reduction in tissues from Tm-treated NSG-MPS I mice compared to untreated NSG-MPS I mice. There was increased, but not significantly different, LAMP-1 staining between the heterozygous and engineered Tm-treated NSG-MPS I mice, supporting the pathologist’s analysis **(Figure 4G)**. Together these data demonstrate that engineered Tm cells were present in several tissues and caused a reduction of GAGs, resulting in attenuated MPS I pathological hallmarks.

### A single dose treatment with IDUA expressing Tm cells histologically improved MPS I pathology but does not significantly improve spatial learning and recognition

Previous studies have shown sustained IDUA levels in the central nervous system leads to improved cognitive function of NSG-MPS I mice.^39–41^ Therefore we investigated the ability of this therapy to improve MPS I hallmarks within the CNS. Immunohistochemical analysis identified human CD3 positive cells with lymphocyte morphology scattered throughout the leptomeninges (indicated by black arrow, bottom inset) **(Figure 5A)**. Human IDUA immunochemistry revealed strong immunopositivity within the choroid plexus in Tm-treated NSG-MPS I mice (middle inset) **(Figure 5A)**. LAMP-1 staining within heterozygous mice showed weak to medium, finely granular cytoplasmic immunopositivity; whereas the untreated and Tm-treated NSG-MPS I mice showed strong, granular cytoplasmic immunopositivity (bottom inset) **(Figure 5A)**. However, quantification of LAMP-1 staining indicated a significant decrease in immunopositivity within the Tm-treated NSG-MPS I mice compared to untreated. Together, these data indicate that there was a slight pathological improvement of the central nervous system in the Tm-treated NSG-MPS I mice.

**Figure 5.**
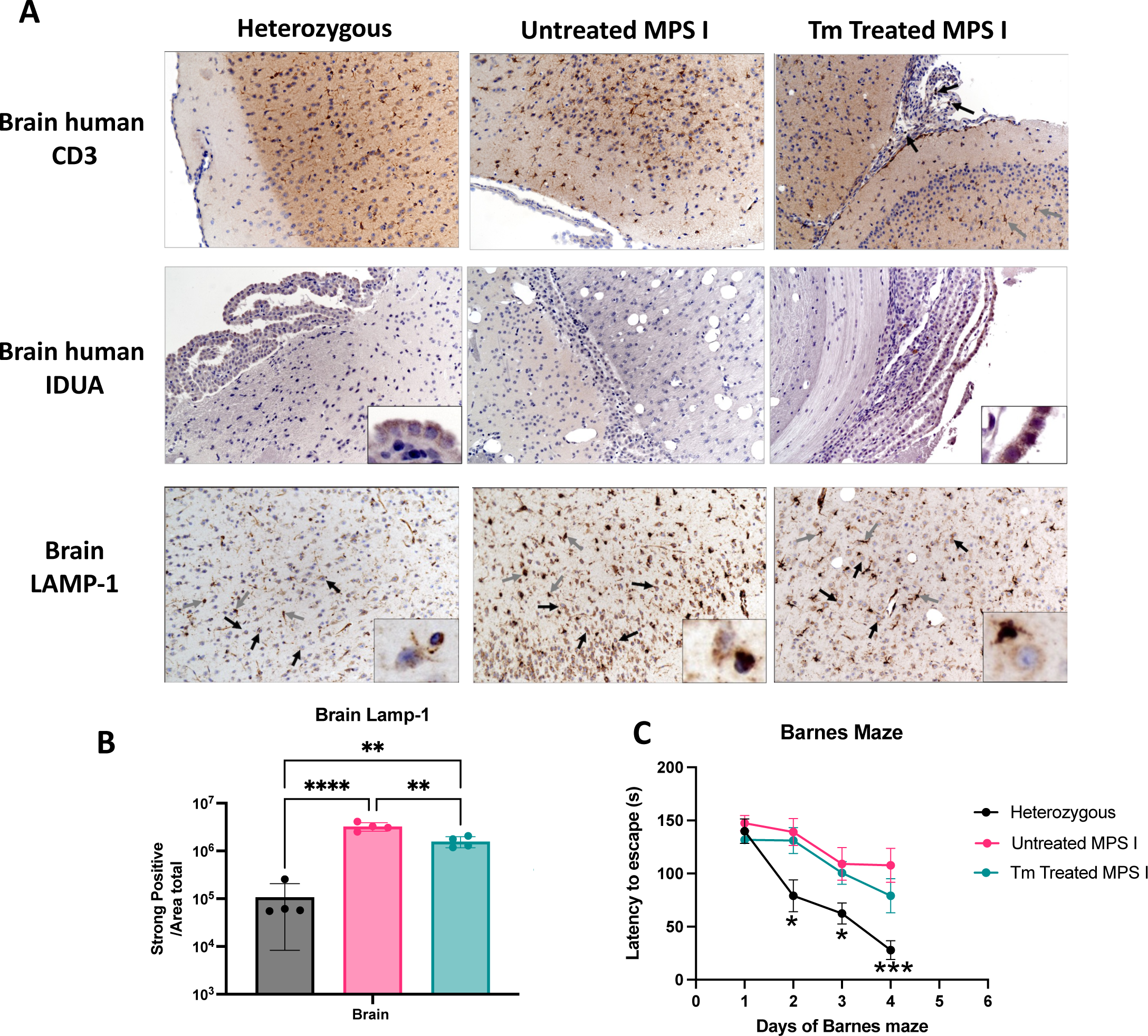
Engineered Tm cells have the capability to ameliorate pathologic MPS I hallmarks within the central nervous system. (A) Representative histological images of human CD3 staining within brain tissue. Black arrows - CD3 immunopositive cells; gray arrows - astrocytes (*top*). Representative histological images of human IDUA within brain tissue (*middle*). Representative histological images of LAMP-1 staining within brain tissue. Black arrows, neurons; gray arrows, astrocytes (*bottom*). (B) Quantification of LAMP-1 immunohistochemistry within brain tissue samples (n=4 each cohort). (C) Barnes maze time to escape of heterozygous (black), untreated NSG-MPS I (red) and Tm-treated NSG-MPS I mice (blue) (n=9). Statistical Analysis: One-way ANOVA (B), Two-way ANOVA (C) *p<0.05, **p<0.01, ***p<0.001, ****p,0.0001.

Lastly, we sought to determine if this therapy is able to measurably impact the neurocognitive function of NSG-MPS I mice. A Barnes Maze assay was used to measure the neurocognitive changes at 20 weeks post-treatment. Over the course of four training days, the heterozygous mice performed significantly better than the other groups **(Figure 5B)**. While the untreated MPS I group’s performance was the worst among all groups, the Tm-treated group had only a slight, non-significant improvement over the untreated MPS I group on day 4 **(Figure 5C)**. This observation indicates that Tm treatment did not significantly improve spatial learning and recognition in the treated mice despite observing decreased LAMP-1 staining and increased IDUA activity in the brain.

## Discussion

Our study explored a novel cell-based therapy using engineered CD4+ Tm cells expressing IDUA to treat a mouse model of MPS I. Employing CRISPR-Cas9 and rAAV6 delivery of donor DNA facilitated high rates of site-specific integration of an IDUA expression cassette into the *AAVS1* locus of Tm cells. These engineered Tm cells were subsequently engrafted into an immunodeficient NSG mouse model of MPS I. Efficacy was assessed by the presence of modified Tm cells, IDUA activity levels, GAG concentrations, and neurocognitive function.

In our short-term study, we demonstrated that engineered Tm cells were able to persist within an NSG murine environment for up to 10 weeks. During this duration, these engineered cells demonstrated the ability to generate and excrete therapeutic IDUA enzyme, enabling systemic metabolic correction in the treated mice. Elevated IDUA within plasma and organ tissue support IDUA cross correction within treated mice. Notably, the secreted IDUA enzyme resulted in systemic reduction of GAG levels across all examined organs and resulted in organ GAG content indistinguishable from heterozygous control NSG mice. However, we observed only a marginal decrease in GAG content in the urine of the cohort at 10 weeks.

In our long-term study, we observed circulating human Tm cells significantly decrease after week 10 in this mouse model, yet a small fraction remained for at least 22 weeks in peripheral blood. Critically, the percentage of engineered Tm cells was maintained at similar levels throughout the study, indicating there is likely not a loss of fitness from the genome engineering process or production of large amounts of IDUA. Initially, the engineered Tm cells led to supraphysiological IDUA activity level at week 6 in peripheral blood, but this activity diminished to a subnormal IDUA activity level after week 12. Tissue GAG content from Tm-treated animals in the 22 week study was increased compared to the 10 week study, yet remained significantly improved in five of the eight organs tested. This may be a result of the Tm cell population decreasing over time after the single dose cell infusion. Moreover, the continual decrease in circulating Tm cell levels at 10 weeks might be linked to inadequate cytokine signaling, or PBMCs within the NSG environment, ultimately negatively impacting the efficacy of the therapy. Overall, the decrease of Tm cells led to reduction of IDUA levels and subsequent tissue GAG content increased significantly from the short-term 10-week study, accumulating in all organs including the brain at 22 weeks post Tm cell treatment. Finally, Barnes Maze assays were utilized to evaluate the spatial learning and cognition of the Tm-treated mice to assess the neurocognitive benefits of our treatment.^42–44^ We noted a subtle cognitive enhancement in the Tm-treated mice, suggesting a potential trend toward neurocognitive improvement. Despite this modest outcome, it’s noteworthy that human IDUA activity was detected across all measured tissues.

A key histopathological hallmark of MPS I, increased LAMP-1 staining, was significantly reduced in liver tissue and to a smaller, yet significant, extent in the brain. This data, combined with the presence of human CD3+ T cells within brain tissue indicates this cell-based therapy has the potential to cross the BBB to correct the neurocognitive manifestations of MPS I. Moreover, sustained IDUA activity led to normalization of tissue GAGs for at least 10 weeks, and urine GAG contents were partially normalized at 22 weeks. However, this therapeutic effect waned over the course of the study, potentially due to the lack of cytokine support or PBMCs in the xenograft model environment. Regardless, improvements for all MSP I manifestations measured were observed at 10 and 22 weeks post treatment administration. Thus, some shortcomings that we encountered in this study need to be addressed in order to improve the overall efficacy of protein secreting, engineered Tm cells.

Numerous strategies can likely be leveraged to enhance the efficacy of Tm cell-based therapies for treating enzymopathies. While our study employed memory CD4+ cells in the artificial NSG model to mitigate graft-versus host disease^45^, research focusing on engineering bulk CD3+ or CD8+ cells could offer a more effective and practical therapeutic avenue as an autologous strategy for human enzymopathy patients. This consideration arises from the fact that CD8+ cells encompass a higher proportion of organ tissue-resident memory cells, especially within the brain, allowing local production and delivery throughout the body’s tissues.^46–49^ Additionally, it may be able to improve performance within this artificial NSG model by providing cytokine support or PBMCs to ensure longevity of Tm cells. Another option would be to administer multiple therapeutic doses to bolster the Tm cell population, which may establish a more enduring therapeutic approach for subsequent investigations. However, these approaches would be addressing an issue with the xenograft model and may not be applicable to an autologous strategy in humans.

In this study we observed minimal albeit detectable levels of Tm cells within the CNS. Unfortunately, this limited infiltration was seemingly inadequate to produce sufficient levels of enzyme in the nervous system, based on a limited effect on neurocognitive function. Instead of solely depending on the infiltration of Tm cells into the brain and CNS, future research might enhance the transport of IDUA enzyme into the CNS via modifications to the enzyme. In fact, ongoing research is exploring the use of fusion IDUA proteins, such as IDUA-antibody fusion proteins or melanotransferrin linked peptide sequence to help shuttle IDUA across the BBB.^38,50,51^ These innovative approaches hold promise for enhancing the delivery of the IDUA enzyme into the brain and CNS, potentially resulting in therapeutic IDUA levels and neurocognitive improvements. Employing these methods in engineered Tm cells would likely offer improved delivery across the BBB as well.

The plasma IDUA levels in our study appeared relatively modest compared to findings from prior research using systemic rAAV or targeted modification of hepatocytes *in vivo* using zinc finger nuclease approaches.^41,52^ Maximizing engineering efficiencies and IDUA production of engineered Tm cells is likely pivotal to producing a viable therapy. Notably, the chromatin state around integrated cassettes can significantly influence transgene expression.^53^ Thus, transgene integration at a more transcriptionally active loci within Tm cells, such as the T cell receptor loci, may hold promise for bolstering IDUA production. Alternatively, employing randomly integrated vectors like transposon and lentiviral delivery methods could amplify engineering efficiencies and IDUA production. However these vectors have oncogenic safety concerts due to their randomly integrating nature.^54,55^ Despite this, studies using lentivirus and transpons to engineer Tm cells for the production of therapeutic proteins have shown great promise.^27,30^ Exploring these engineering alternatives may provide a way to create a more effective cell-based therapy.

Expanding upon these findings, it will be imperative to investigate IDUA production from engineered Tm cells within an immunocompetent setting using murine Tm cells. This would allow for the investigation of the host immune response to IDUA and the engineered Tm cells that an NSG mouse model cannot address. As Newborn screening for MPS I is now recommended in the US this enables earlier initiation of ERT.^56,57^ In-utero ERT administration has demonstrated increased immune tolerance towards lysosomal enzymes.^58^ Consequently, the likelihood of immune clearance of the engineered cells producing IDUA is reduced. Moreover, T cells are not professional antigen presenting cells and may thus be less likely to be cleared from the body due to expression and presentation of IDUA derived antigens.^59^ Therefore, exploring the use of engineered murine Tm cells for adoptive transfer into an immunocompetent MPS I model would represent additional studies to further support or refute the use of engineered Tm cells for enzyme replacement therapy. This approach could facilitate prolonged engraftment and sustained therapy, providing valuable insights for long-term therapeutics strategies.

Engineering antigen-specific T cell populations or delivering a known T cell receptor (TCR) may provide an avenue for *in vivo* T cell boosts, therefore increasing engineered cell populations and IDUA levels. Epstein-Barr virus (EBV) antigen was previously used to boost a murine T cell adoptive therapy *in vivo* for the delivery of erythropoietin to treat anemia.^30^ Moreover, workflows for isolating and expansion of various antigen specific T cells are well established in the cancer field.^60,61^ An available population of antigen-specific memory CD4+ and CD8+ T cells are developed from routine childhood vaccination, such as TDaP vaccination.^62–64^ Such vaccinations give rise to memory T cell populations that not only endure for years, but can also be reactivated upon subsequent exposure to the corresponding pathogen or vaccine.^65,66^ Harnessing these antigen-specific memory T cells could result in a boostable cell-based therapy to supply IDUA, potentially creating a lifelong therapeutic platform derived from peripheral blood samples early in life.

Allogeneic HSCT and ERT have increased the quality of life for many patients with Hurler syndrome. However, many difficulties remain; particularly the fact that the neurocognitive benefits are limited with these treatments. Other residual disease burdens persist as well, such as non-progressive mental deficits, orthopedic manifestations, and damage to various organs.^67–69^ Allogeneic HSCT is considered the most effective treatment for MPS I patients under two years of age; however, the long-term effects of HSCT can lead to an increased risk of life-threatening infections, as well as a substantial risk for GVHD.^70^ A therapy using autologous HSPCs transduced with an IDUA-encoding lentiviral vector is currently under clinical investigation with promising results.^71^ However, lentiviral transduction is not risk free, with the potential of insertional mutagenesis leading to transformation.^72^ Further, HSPC transplant conditioning treatments are not risk free.^73^ Various *in vivo* rAAV delivery strategies are under exploration for therapeutic protein delivery as well.^74–76^ rAAV has been used to deliver transgenes for treatment of MPS I and MPS II into the central nervous system and currently there is an ongoing clinical trial using rAAV9 to deliver IDUA into the central nervous system of patients (NCT03580083).^39,77^ However, meta-analysis of 255 rAAV clinical trials calls into question the cost, durability, immunogenicity, and toxicity of various *in vivo* rAAV gene therapies.^78^

Utilizing autologous engineered Tm cells reduces many of the aforementioned risks of exploratory therapies for enzymopathies, making our approach more attractive than *in vivo* gene delivery or use of engineered autologous HSPCs. Notably, autologous Tm cells are easily obtained from peripheral blood, can be re-dosed, are long lived, are unlikely to induce GVHD, have the potential for antigen-specific boosts, and are capable of taking permanent residence in all tissues, including in the brain and CNS.^14,79^ Our proposed cell-based therapy also benefits from an established clinical and manufacturing pipeline, pioneered by the CAR-T industry. Thus, IDUA expressing Tm cells may offer a unique and distinct advantage over current therapies and other exploratory approaches for the treatment of MPS I.

Our study demonstrates the potential use of engineered Tm cells for cellular-based gene therapy of MPS I. We observed IDUA activity in plasma and tissues of treated mice that resulted in normalization of the tissue pathology and GAG levels after a single dose of engineered Tm cells. Although we observed limited benefits of the treatment to neurocognitive function, this study demonstrates that engineered Tm cells have the potential for use in a cellular-based gene therapy for MPS I and may be applicable for other enzymopathies as well. Our research, in conjunction with previous studies,^27,30^ provides further evidence that T cells are a promising platform for delivery of therapeutic enzymes for the treatment of enzymopathies.

## Materials and Methods

### T cell isolation

Human leukapheresis samples from de-identified, healthy donors were obtained by automated leukapheresis (Memorial Blood Centers, Minneapolis, MN) with approval from the University of Minnesota Institutional Review Board (IRB 1602E84302). Leukapheresis samples were processed with a Ficoll-paque (GE Healthcare Life Sciences) density gradient in accordance with the manufacturer’s instructions. Cells were then resuspended in ammonium-chloride-potassium (ACK) lysis buffer (Quality Biological) and incubated for three minutes at room temperature (RT) to lyse red blood cells. CD4^+^ CD45RO^+^ memory T cells were then isolated by negative selection using an EasySep^TM^ Human Memory CD4+ T Cell Enrichment Kit (STEMCELL Technologies).

### T cell recovery medium

Optimizer^TM^ T-Cell expansion basal medium (Gibco) was supplemented with 2.6% OpTmizer^TM^ T-Cell expansion supplement (Gibco), 2.5% CTS^TM^ Immune cell serum replacement (Gibco), 1% L-Glutamine (Gibco), 10 mM N-Acetyl-L-cysteine (Sigma-Aldrich) and pre-equilibrated to 37°C prior to use.

### T cell culture medium (TCM)

As previously described,^19^ Optimizer^TM^ T-Cell expansion basal medium (Gibco) was supplemented with 2.6% OpTmizer^TM^ T-Cell expansion supplement (Gibco), 2.5% CTS^TM^ Immune cell serum replacement (Gibco), 1% L-Glutamine (Gibco), 1% Pencicillin/Streptomycin (Millipore), 10 mM N-Acetyl-L-cysteine (Sigma-Aldrich), 300 IU/ml recombinant human IL-2 (Prepotech), 5 ng/ml recombinant human IL-7 (Prepotech), 5 ng/mL recombinant human IL-15 (Prepotech) and pre-equilibrated to 37°C prior to use.

### T cell stimulation

As previously described,^19^ isolated memory CD4^+^ T cells (Tm) were plated in a well of 24-well plate containing TCM at a density of 1 x 10^6^ cells/mL. Research-grade Dynabeads^TM^ Human T-Expander CD3/CD28 beads (Gibco) were washed using DynaMag-2 magnet (Gibco) once with 1xPBS and added to the cells at 2:1 ratio (beads:cells). The plate was incubated at 37°C, 5% CO_2_, with humidity for 48 hours.

### Recombinant AAV pseudotyped 6 (rAAV6) vector

rAAV6 AAVS1 SA-STOP-pA-MND-IDUA-RQR8 was used as a DNA donor template for homology-directed repair (HDR). The vector construct is comprised of homology arms targeting the AAVS1 locus of the Tm cells and consists of an MND promoter, the IDUA-T2A-RQR8 sequence, and a BGH poly-A-tail. The construct was cloned into an AAV2 backbone and packaged using AAV6 capsid at Vigene Biosciences.

### CD4+ cell engineering and expansion

Dynabeads were dislodged from the stimulated Tm cells by pipette mixing and then separated from the Tm cells using a DynaMag-2 magnet (Gibco). Tm cells were washed once with 1X PBS with centrifugation at 200 x g for 10 minutes. The cells were resuspended in 20 μL P3 Nucleofector Reagent (NFR) per manufacturer’s instructions (P3 Primary cell 4D-Nucleofector, Lonza). Transfection reaction was prepared by adding 1.5 μg chemically modified SpCas9 mRNA (CleanCap^®^, Trilink) along with 1 μg chemically modified sgRNA (Integrated DNA Technologies) targeting the *AAVS1* locus into the cell suspension. The transfection reaction was transferred into P3 Primary cell cuvette and electroporated with EO-115 program on a Lonza 4D Nucleofector (Lonza). The cells were rested in the cuvette without disturbing them for 15 minutes at RT. Then the cells were transferred into a well of a 24-well plate containing 300 μL T cell recovery media for additional 30 minutes at 37°C and 5% CO_2_. rAAV6 SA-STOP-pA-MND-IDUA-RQR8 locus was then added to the resting cells at an MOI of 500,000. Then 700 μL of TCM containing Dynabeads at a ratio 1:2 (beads:cells) were immediately added to the cells. Cells were then incubated at 37°C and 5% CO_2_ for an additional 24 hours.

After 24 hours, the Tm cells were moved to a well of a 24-well G-Rex tissue culture plate (Wilson Wolf) and brought up to 6.6 mL with TCM. The engineered Tm cells were cultured for an additional seven days. Half media changes in the G-Rex with TCM with 2X cytokine supplement were made every three days without disturbing the cells.

### Animal care and husbandry

All animal care and experimental procedures were conducted under the approval of the Institutional Animal Care and Use Committee (IACUC) of the University of Minnesota. Transgenic mice heterozygous for an IDUA knockout allele on the NOD.Cg-Prkdc^scid^Il2rg^tm1Wjl^/SzJ background (NSG-IDUA^+/-^, heterozygous) breeding pairs were kindly provided as breeding pairs by Dr. Schwartz (Children’s Hospital of Orange County, CA)^31^ and were maintained under specific pathogen free conditions at the Research Animal Resources (RAR) facility of the University of Minnesota. PCR amplification with a mixture of a common forward primer (5’-GGAACTTTGAGACTTGGAATGAACCAG-3’), a WT reverse primer (5’-CATTGTAAATAGGGGTATCCTTGAACTC-3’) and a mutant reverse primer (5’-GGATTGGGAAGACAA TAGCAGGCATGCT-3’) were used to amplified WT or mutant allele, as described previously (PMID: 26052536).

### Engraftment of IDUA-expressing Tm cells into NSG-MPS I mice

Two studies were carried out: a short-term study (10 weeks) and a long-term study (22 weeks). Three groups of mice were used for each study: heterozygous littermate control, untreated littermate MPS I control, and Tm-treated MPS I (short-term study n=6 per group,long-term study n=12 per group). At three weeks old all mice were pre-conditioned for engraftment with 25 mg/kg busulfan (Millipore-Sigma) via intraperitoneal injection (IP). 48 hours later, Tm-treated NSG-MPS I mice received 1x10^7^ unsorted engineered Tm cells in 200 μL sterile 1x PBS via IP injection. The heterozygous and untreated NSG-MPS I mice received 200 μL of sterile 1xPBS via IP injection at the engraftment day (Supplementary Figure 2).

### Blood and plasma collection

Mice were bled and blood was collected in a K3-EDTA tube (Greiner Bio-One) two days prior to engraftment and every two weeks post engraftment throughout the study. Blood was centrifuged at 550 x g for 8 minutes at 4°C and the plasma fraction was collected for use in the IDUA assay. ACK lysis buffer (Theromo Fisher) was added to the blood cell fraction on ice for twenty minutes and then washed in cold dPBS (Theromo Fisher). Cells were then strained through a 70µm filter and analyzed by flow cytometry.

### Urine collection

A total of 100-200 μl of urine was collected by gentle massage of the bladder around the abdominal area of the mice at day -2 prior to engraftment and then every six weeks post-engraftment through the end of the study. The urine was centrifuged at 12,000 x g for 5 minutes to pellet any debris before use.

### Organ collection

Mice were sacrificed and immediately perfused by hand pressure with 0.89% normal saline (sodium chloride, Acros Organics) using an infusion wing set with 23GX3/4 gauge needle with 12” tubing (Terumo). The heart, lung, liver, spleen, kidney, spinal cord, brain, and bone marrow were collected and divided into three sections: one third of each organ was processed using a plastic pestle (Corning) to press the organ against a 70 micron cell strainer (Falcon), and the other part of each organ was placed in a clean 1.5 mL microcentrifuge tube with 200-500 mL of ice-cold saline and homogenized with FisherBrand Sonicator model FB50 with probe CL-18 (Fisher Scientific) settings of 30 Amplitude for 10 seconds for a total of 3 intervals (on ice). The homogenates were centrifuged at 13,000 x g for 25 minutes at 4°C. The tissue lysates were then transferred into a new 1.5 mL microcentrifuge tube and stored at -80°C until use. The remaining section of each tissue was fixed in 10% buffered formalin for 24 hours then washed in 100% ethanol before being sent for histological analysis.

### Flow cytometry analysis

Cells from processed organ and blood samples are washed twice with flow buffer (1x PBS (Gibco), 0.5% BSA (Sigma-Aldrich), 2 mM EDTA (Invitrogen) and pre-treated with a mixture of human TruStain FcX (Fc Receptor blocking solution, Biolegend) and CD16/CD32 Rat anti-mouse Fc antibodies (mouse BD Fc blocker, BD Biosciences) for 5 minutes at RT and washed twice with flow buffer. Cell pellets were then incubated with Fixable Viability Dye eFluor780 (eBioscience) for 10 minutes at RT. Cells were then washed once with flow buffer and labeled with fluorophore-conjugated antibodies (see supplementary table 1) for 15 minutes in the dark at RT. Cells were then washed once and resuspended with 1% paraformaldehyde and analyzed on a CytoFlex S (Beckman Coulter) flow cytometer. Data analysis was performed with FlowJo version 10.7.1 software (BD Biosciences).

### IDUA activity assay

As previously described, 4-methylumbelliferyl α-L-iduronide (4-MU-iduronide, Glycosynth) IDUA enzyme substrate was diluted in 0.4M sodium formate buffer at pH 3.5 to make 360 μM working solution.^32^ Twenty-five microliters of the working substrate solution was mixed with 25μl of sample (i.e. mouse plasma, the culture medium of engineered T cells, or tissue lysate) and incubated at 37°C for 30 minutes. Then 200 μl of stop buffer (Glycine carbonate buffer, pH 10.4) was added to the sample to stop the enzyme activity. 4-methylumbelliferone (4-MU) was prepared in different dilutions and used to make a standard curve. Fluorescence was measured using a Bio-Tek plate reader with excitation at 355 nm and emission at 460 nm. IDUA enzyme activity was calculated from fluorescence intensity by using a standard curve. Plasma or culture media IDUA activities are reported as nmol/hour/ml of the respective sample. Tissue IDUA activities are reported as nmol/hour/mg protein of the respective sample. Protein mass was determined using the BCA protein assay kit (Abcam) with PierceTM bovine serum albumin as a standard (Thermo Scientific).

### GAGs assay

Tissue lysates were incubated overnight at 55°C with Proteinase K, DNase1, and RNase. GAG contents were assessed using the Blyscan™ Sulfated Glycosaminoglycan kit (Biocolor Life Science; Accurate Chemical) in accordance with the manufacturer’s instructions. Blyscan GAG standard 100 μg/ml (Accurate Chemical) was used to generate a standard curve. The end-product was measured for absorbance at 656 nm using Synergy BioTek plate reader and Aglienet Gen5 program, version 3.14. Tissue GAG content is reported as μg/mg protein. Urine GAG was assessed without processing of the urine. Urine GAG is reported as μg/mg creatinine. A Creatinine Assay Kit (Sigma-Aldrich) was used to measure creatinine in the urine in accordance with the manufacturer’s instructions.

### Barnes maze

The Barnes maze with AnyMaze (San Diego Instrument) program, version 7.0, was used to assess spatial learning and memory. In brief, the Barnes maze is an elevated circular platform with a total of 20 holes that are evenly shaped and spaced at the periphery. All the holes were blocked except one hole with an escape box. The escape box was maintained at the same position of the room throughout the 4-day course of experiment. Four spatial cues were placed on each of the four walls, and the room was lit with 300 lux. Mice were acclimated in the dark for at least 15 minutes at the beginning of each training session. The mouse was placed at the center of the platform with a dark cover, then the light was turned on and the cover was removed simultaneously to allow the mouse to explore under the bright light. The animal was expected to escape the platform using spatial navigation to the open hole with the escape box within three minutes. If the mouse was unable to find the exit by the end of 3 minutes, it was gently guided to the hole, which was subsequently closed. Each mouse was subjected to four trials a day for a total of four days. The latency to escape was recorded for each trial, and the average latency to escape was calculated for each day in each group.

### Histology

Immunohistochemical staining and analysis was completed by the Comparative Pathology Shared Resource at the University of Minnesota. Tissues were formalin-fixed and processed through a series of graded alcohols, embedded in paraffin, and individual sections were cut and stained on individual mounted slides. H&E-stained sections were evaluated using light microscopy. Prepared tissue sections were deparaffinized, rehydrated, and then stained with rabbit monoclonal anti-LAMP antibody (ab208943; Abcam), anti-human CD3 epsilon antibody (MAB10670; R&D Systems), or anti-human alpha-L-Iduronidase antibody (AF4119; Biotechne). Quantification and analysis was performed with Aperio Imagescope software for human IDUA and LAMP1 image sets. Algorithm Positive Pixel Count v9, default settings except Ip(Low) = Isp(High) set to 75 for liver samples and Ip(Low) = Isp(High) set to 50 for brain samples.

### Statistical analysis

The Student’s t-test was used to evaluate the significant differences between two conditions. Differences between the means of three or more experimental groups were evaluated by a one-way ANOVA test. Differences between three or more experimental groups with multiple data points were evaluated by a two-way ANOVA test. Means values + SD are shown for all results. The level of significance was set at p<0.05. All statistical analyses were performed using GraphPad Prism 9.2.0 (GraphPad).

## Data Availability

All reasonable requests can be made to the corresponding author at

## Acknowledgements

B.R.W. acknowledges funding from Office of Discovery and Translation, NIH grants R21CA237789, R21AI163731, P01CA254849, P50CA136393, U54CA268069, R01AI146009, and, Children’s Cancer Research Fund. B.S.M acknowledges funding from NIH grants R01AI146009, R01AI161017, P01CA254849, P50CA136393, U24OD026641, U54CA232561, P30CA077598, U54CA268069, Children’s Cancer Research Fund, the Fanconi Anemia Research Fund, and the Randy Shaver Cancer and Community Fund. We would like to thank Philip H. Schwartz for providing the NSG-MPS I mice.

## Author Contributions

K.L. designed experiments. K.L. E.W.K., and J.D.J. performed assays, analyzed the data, and wrote the manuscript. J.J.P. assisted with experimental assays. N.J.S., B.J.W., M.J.J., assisted with technical expertise. B.R.W. and B.S.M supervised the research.

