## Supplemental Figures for "Engineering Memory T Cells as a platform for Long-Term Enzyme Replacement Therapy in Lysosomal Storage Disorders"

**Supplementary Figures**

Supplementary table 1

Supplementary table 1. Antibodies for flow cytometry.

| Antibody | Fluorophore | Clone | Catalog no. | Company |
|---|---|---|---|---|
| Human CD45 | AF700 | 2D1 | 368514 | Biolegend |
| Mouse CD45 | APC | 30-F11 | 103112 | Biolegend |
| Human CD45RO | Cy7PE | UCHL1 | 304230 | Biolegend |
| Anti-CD34 (RQR8) | PE | QBEND/10 | MA1-10205 | ThermoFisher |

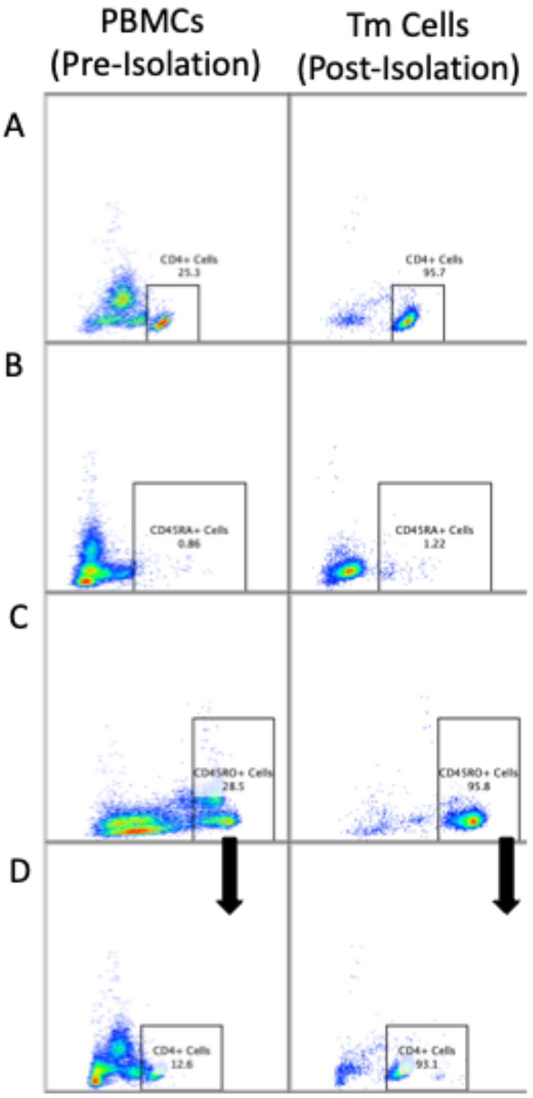


Figure S1

Flow cytometry plots of isolated donor memory T cells prior to engineering and delivery into NSG-MPS I mice. (A) CD4 staining of peripheral blood mononuclear cells (PBMCs) before Tm isolation and post Tm isolation. (B) CD45RA staining of peripheral blood mononuclear cells

(PBMCs) before Tm isolation and post Tm isolation. (C) CD45RO staining of peripheral blood mononuclear cells (PBMCs) before Tm isolation and post Tm isolation. (D) CD4+ population of the CD45RO+ population of peripheral blood mononuclear cells (PBMCs) before Tm isolation and post Tm isolation.

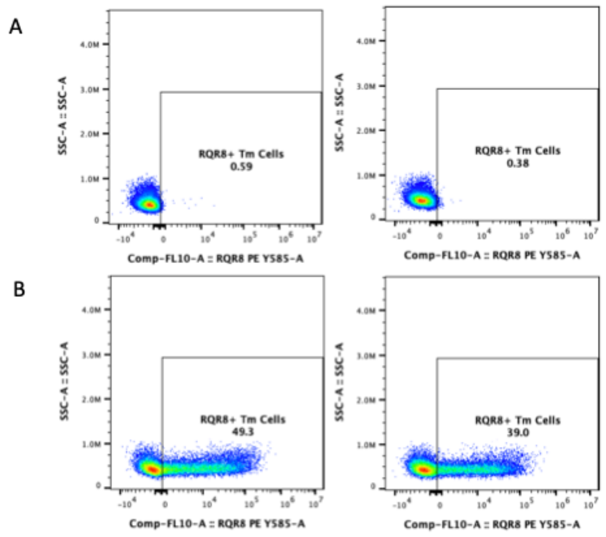


Figure S2

Flow cytometry plot of RQR8 expression in Tm cells from two donors 11 days post engineering with the rAAV containing IDUA transgene. (A) Vector controls cells (B) rAAV engineered cells.

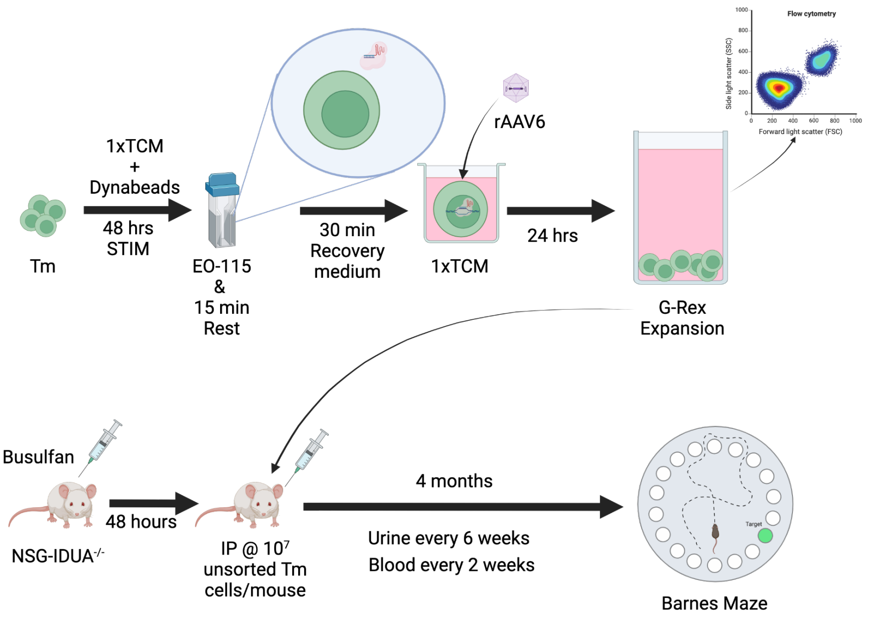


Figure S3

Graphical outline of engineering and subsequent engraftment of Tm cells. Enriched Tm cells are activated for 48 hours and are electroporated delivering delivery gene editing reagents. Donor DNA is delivered using rAAV6 30 minutes post electroporation. Cells are then expanded and engineered efficiency is measured through flow cytometry. NSG mice are preconditions for cell delivery with busulfan 48 hours before conditions with busulfan 48 hours before cell injections. Mice were followed for 22 weeks with periodic blood and urine samples taken. Neurocognitive tests were performed with Barnes maze at the end of the 22 week time. Mice were then sacrificed for tissue analysis.

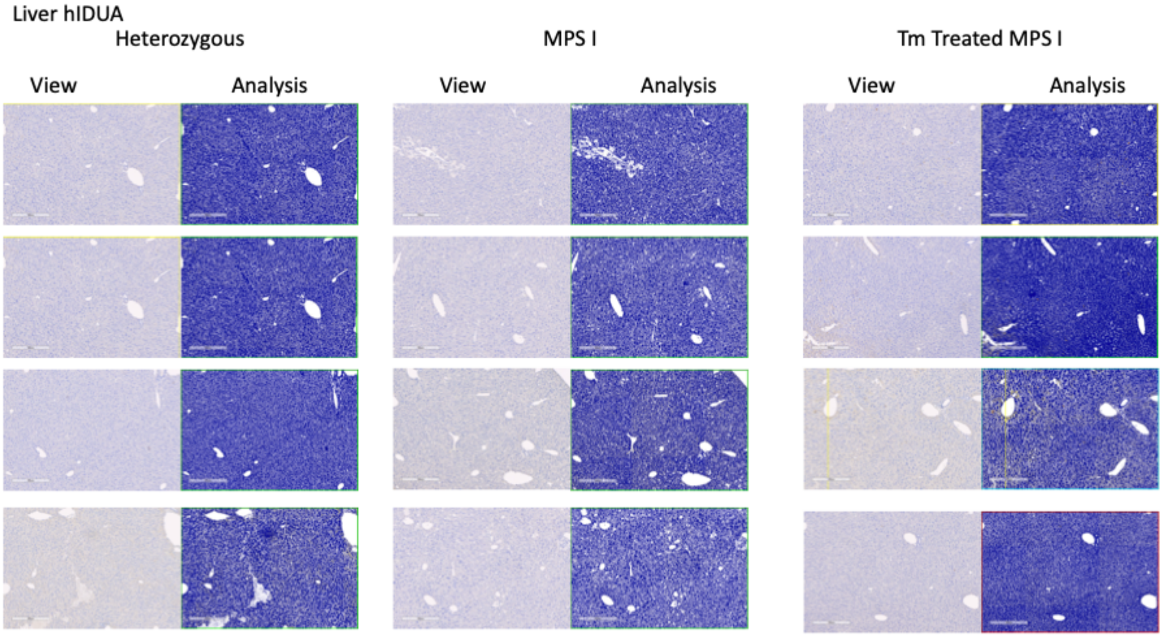
Liver hIDUA
Heterozygous
MPS I
Tm Treated MPS I
View
Analysis
View
Analysis
View
Analysis

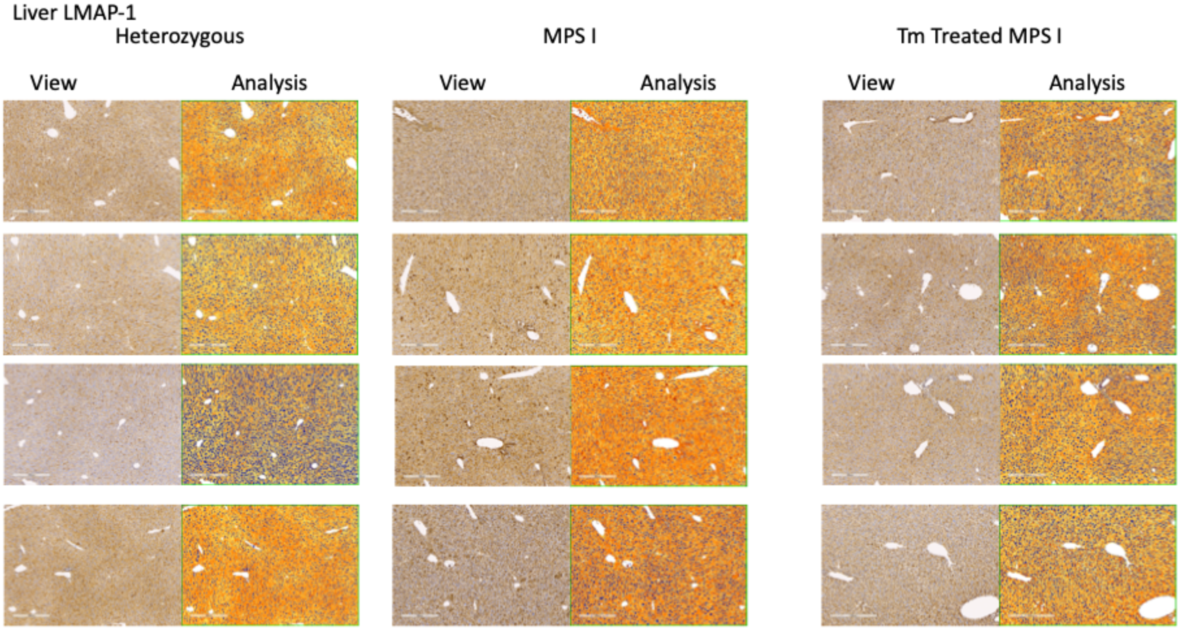
Liver LMAP-1
Heterozygous
MPS I
Tm Treated MPS I
View
Analysis
View
Analysis
View
Analysis

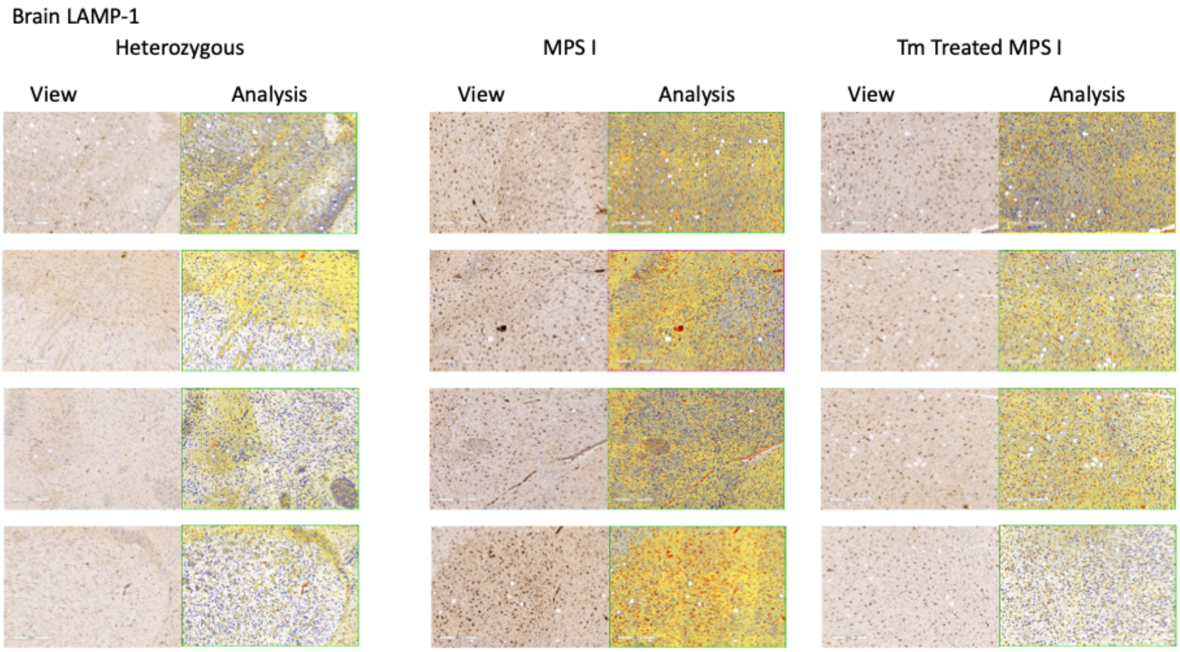


Figure S4

Immunohistochemistry image analysis. Raw images (left side of columns) and analyzed images (right side of columns) used for analysis of human IDUA and LAMP-1 quantification. n=4 each for heterozygous, untreated NSG-MPS I, and Tm-treated NSG-MPS I mice.
